# Hepatic NFAT signaling regulates the expression of inflammatory cytokines in cholestasis

**DOI:** 10.1101/2020.03.25.007831

**Authors:** Shi-Ying Cai, Dongke Yu, Carol J. Soroka, Jing Wang, James L. Boyer

## Abstract

The inflammatory response plays an important role in cholestatic liver injury where bile acid (BA) induction of proinflammatory cytokines in hepatocytes is an initial pathophysiologic event. However, the signaling pathways involving BA stimulation of cytokine production remain to be elucidated. In this report, we examined the functional role of the Nuclear Factor of Activated T-cells (NFAT) in BA-induction of inflammatory genes in hepatic cells and cholestatic livers. We found that NFAT isoform c1 and c3 were expressed in human and mouse hepatocytes. When treated with cholestatic levels of BA, both human and mouse hepatocytes but not cholangiocytes increased NFATc3 nuclear translocation, associated with elevated mRNA levels of IL-8, Cxcl2, and Cxcl10 in these cells. Blocking NFAT activation with pathway-specific inhibitors (i.e. cyclosporine A, FK-506, KN-62 and Inca-6) or knocking down Nfatc3, significantly repressed BA-induction of these cytokines in mouse hepatocytes, including Ccl2, Cxcl2, Cxcl10, Icam1 and Egr1. Nuclear expression of NFATc3/Nfatc3 protein was also increased in cholestatic livers after bile duct ligation or in Abcb4^-/-^ mice and in patients with primary biliary cholangitis and primary sclerosing cholangitis in association with tissue elevations of Cxcl2 and IL-8. Gene reporter assays and ChIP-PCR demonstrated that the NFAT response element in its promoter played a key role in BA-induced human IL-8 expression. Together our findings indicate that NFAT plays an important role in BA stimulation of hepatic cytokines in cholestasis and is a mechanism that may provide novel targets to reduce cholestatic liver injury.

## Introduction

Cholestasis is a syndrome where bile acids (BA) accumulate in the liver, resulting in liver injury. Cholestasis can be caused by genetic or developmental defects, as well as result from acquired diseases (1-3). In many cases, including primary biliary cholangitis (PBC), primary sclerosing cholangitis (PSC), and biliary atresia, the etiology is not known. Yet regardless of the initial cause, elevated hepatic BA concentrations are common to all. However, the pathogenetic role of BA in cholestasis remains unclear, hindering the development of effective therapies for these disorders. Recent studies indicate that the inflammatory response plays an important role in the pathogenesis of this injury and that BA induction of proinflammatory cytokines in hepatocytes is the initial pathophysiologic event (4-6). In the process of this event, BA cause endoplasmic reticulum (ER) stress and mitochondrial damage in hepatocytes. These injured mitochondria release DNA that then activates the Toll-like receptor (TLR/Tlr) 9 signaling pathway that results in the up-regulation of inflammatory cytokine expression(4). However, hepatic specific deficiency of Tlr9 diminishes, but does not eliminate, cholestatic liver injury in mice, indicating that signaling pathways independent of Tlr9 must also play a role in its pathogenesis (4). During these investigations we found that cyclosporine A (CsA)(4), a potent inhibitor of the calcium (Ca^2+^) / calcineurin / Nuclear Factor of Activated T-cells (NFAT) signaling pathway, significantly repressed BA induction of inflammatory cytokines in mouse hepatocytes, suggesting that the transcription factor NFAT is activated in cholestatic hepatocytes. NFAT is a highly phosphorylated family of proteins, described initially in T-cells (7-10). There are four isoforms of NFAT that are activated by intracellular Ca^2+^ signaling, i.e. NFATc1, c2, c3 and c4. These isoforms are differentially expressed in tissues and cells, and are activated in a stimulus-dependent and isoform-specific manner(11-13). At resting state, NFAT is localized in the cytosol. However, when there is a sustained elevation of intracellular [Ca^2+^], NFATc is dephosphorylated and translocates into the nucleus. In the nucleus NFATc regulates the expression of its targets either by binding directly to its specific response elements or by associating with other transcription factors on the promoter of the target. Many of NFAT targets in T-cells are involved in the immune response(14). However, it is not known whether NFAT plays any role in the BA-induced hepatic inflammatory response.

In this report, we examined the functional role of NFATc3 in cholestatic liver injury in mice and humans. We found that Nfatc3 mediated BA induction of inflammatory cytokines in mouse hepatocyte cultures. Knockdown of Nfatc3 reduced BA induction of these cytokines. Increased nuclear expression of NFATc3/Nfatc3 protein was also seen in the livers of patients with PBC and PSC as well as *Abcb4*^-/-^ mice, associated with elevated tissue levels of IL-8 and Cxcl2 in humans and mice, respectively. BA-stimulated IL-8 expression is mediated through an NFAT response element in the human IL-8 promoter as determined by gene reporter assays. Together, these findings describe a previously unrecognized signaling pathway in hepatocytes that contributes to BA initiated hepatic inflammation, providing novel strategies for treating cholestatic liver diseases.

## Materials and Methods

### Materials

Chemicals were purchased from Sigma-Aldrich (St. Louis, MO), except where otherwise specified. Cell culture media (DMEM and Williams’ E), fetal bovine serum (FBS), penicillin/streptomycin, trypsin, phosphate buffered saline (PBS), Lipofectamine 2000 and Lipofectamine RNAiMax were from Life Technologies (Carlsbad, CA). Collagen coated plates and collagen were purchased from BD Sciences (Bedford, MA). Mouse Nfatc3 siRNA oligoes (siGENOME SMARTpool) and control siRNAs were purchase from Dharmacon (Lafayette, CO). FK506 was purchased from Cayman Chemical (Ann Arbor, MI). KN-62 and Inca-6 were from Tocris Bioscience (Minneapolis, MN). ECL reagents were from Thermo Scientific.

### Preparation and maintenance of mouse hepatocytes, cholangiocytes and human hepatocytes

Mouse hepatocytes were isolated from 10-20 weeks old C57Bl/6 mice using collagenase perfusion. Primary cultures of mouse cholangiocyte were provided by Dr. Strazzabosco’s lab here in our Liver Center(15). Human hepatocytes were obtained through the Liver Tissue Cell Distribution System (Pittsburgh, Pennsylvania), which was funded by NIH Contract #N01-DK-7-0004 / HHSN267200700004C. Both human and mouse hepatocytes were maintained as previously described(16). All cell cultures were treated with indicated chemicals and collected within 96 hr after isolation. Protein and mRNA expression were detected as described(17). Antibodies against NFATc3 were purchased from Santa Cruz Biotechnology (Cat# sc-8405), GAPDH antibody was from Sigma-Aldrich, Lamin B1 antibody (Cat#12586S) was from Cell Signaling Technology (Danvers, MA), and Nucleoporin p62 antibody (Cat#51-9002029) was from BD Biosciences. To transfect mouse hepatocytes, siRNAs were mixed with Lipofectamine RNAiMax in Opti-MEM (Life Technologies) following manufacturer’s instruction and added to the culture medium 3 hr after the cells were plated on collagen coated plates (4×10^5^ cell/well in a 12-well plate). Forty hours after transfection, the cells were treated with BA for 24 h, and collected for gene expression analyses.

### Preparation of nuclear and cytoplasmic proteins from cells and tissues

Frozen mouse liver was available from our previous studies(4, 18). De-identified human liver tissue specimens came from Yale Liver Center and the Liver Tissue Cell Distribution System in University of Minnesota, Minneapolis, Minnesota, which was funded by NIH Contract # HSN276201200017C. Cholestatic liver specimens were explanted from patients with end-stage PBS or PSC, while normal human liver tissues were from deceased donors who did not have liver diseases. To isolate cytoplasmic and nuclear protein from cells and tissue, a kit from Pierce Biotechnology (Cat# 78833, Rockford, IL) was used following manufacturer’s instructions. A protease inhibitor cocktail (Cat# 11873580001, Roche Diagnostic GmbH) was added to the solutions of the preparation. A dounce homogenizer was used to gently break up the liver tissue. In PBS. The homogenate was centrifugated at 500x*g* for 3 min, and the pellet was used as the starting material.

### Liver immunohistochemistry (IHC)

Formalin fixed, paraffin-embedded mouse liver was tested for Nfatc3 localization using mouse anti-Nfatc3 antibody (Santa Cruz Biotechnology, Cat# sc-8405) and the NovoLink Polymer Detection System (Cat# RE7140-K) from Leica Biosystems (Newcastle, UK) according to the manufacturer’s instructions. Antigen retrieval was achieved with citrate buffer at pH 6.0 in a steamer for 20 min and primary antibody incubated on the tissue section overnight at 4°C. Sections were counter-stained with hematoxylin.

### Human IL-8 promoter reporter assay and Chromatin Immunoprecipitation (ChIP)-PCR assay

The proximal promoter region of human IL-8 gene was amplified from a BAC clone (CH17-113M16, purchased from Children’s Hospital Oakland Research Institute, CA) using PCR with Forward primer: GCATGGTACCAG ATCTTCACCATCATGATAGCATCTGTA and Reverse primer: GCATGGATCCTGGCTCTTGTCCTAGAAGC. The amplified fragment (490 bp) was cloned into pGL3-basic vector, and verified by DNA-sequencing. The NFAT response element in human IL-8 promoter was mutated from GGAATT**TCC**TCT to GGAATT**CTT**TCT. For gene reporter assay, the reporter constructs were transfected into Huh7-BAT cells(19) (a cell line stably transfected with human NTCP, provided by Dr. Gregory Gores, Mayo Clinic) using Lipofectamine 2000 by following the manufacturer’s instruction. Twenty-four hours after transfection, the cells were treated with indicated concentration of BA for 16 hr. Dual-luciferase assay (purchased from Promega) were performed to determine promoter activity. Data are normalized to Renilla luciferase activity. For ChIP-PCR, Huh7-BAT cells were treated with indicated BA for 4 hr. After the cells were cross-linked, the assay was performed using a kit from Pierce Biotechnology (Pierce Agarose ChIP Kit, Cat#26156) by following the manufacturer’s instruction. Specifically, for each immunoprecipitation reaction, the nuclear fraction from 2×10^6^ cells and 2 µg of antibody or control IgG were used. The primers for the detection PCR are TGGGCCATCAGTTGCAAATCG as the forward primer and the above-mentioned Reverse primer as the reverse primer. It generates a 170bp band.

### Statistical Analysis

One-way ANOVA followed by student *t* test were used to perform the statistical analysis for liver tissues. Paired t-test was used for dual-luciferase reporter assays. Data are presented as the means ± S.D. *P<*0.05 was considered statistically significant.

## Results

### Inhibitors involved in NFAT signaling repressed BA induction of chemokines in mouse hepatocytes

To confirm that CsA specifically blocks BA induction of chemokines in mouse hepatocytes as we described previously (4), different doses of CsA were added to mouse hepatocytes exposed to 50 μM glycocholic acid (GCA) for 24 h (Fig 1). CsA demonstrated dose-dependent inhibition of GCA induction of both Cxcl2 and Cxcl10 in these cells with the optimal concentration at 3 µM. To examine whether other inhibitors involved in Ca^2+^-calmodulin-calcineurin-NFAT signaling pathway could also block BA induction of these chemokines (Figure 1B), we treated mouse hepatocytes exposed to 50 μM GCA with KN-62 (a specific Ca^2+^/calmodulin-dependent protein kinase inhibitor), FK506 (which binds to FKBP12 and blocks calcineurin activation) and Inca-6 (an inhibitor that prevents calcineurin and NFAT binding). Each of these inhibitors significantly reduced GCA induction of Cxcl2 (Fig 1C). Together, these findings indicate that Ca^2+^/NFAT signaling pathway is involved in BA induction of chemokines in mouse hepatocytes.

**Figure 1.**
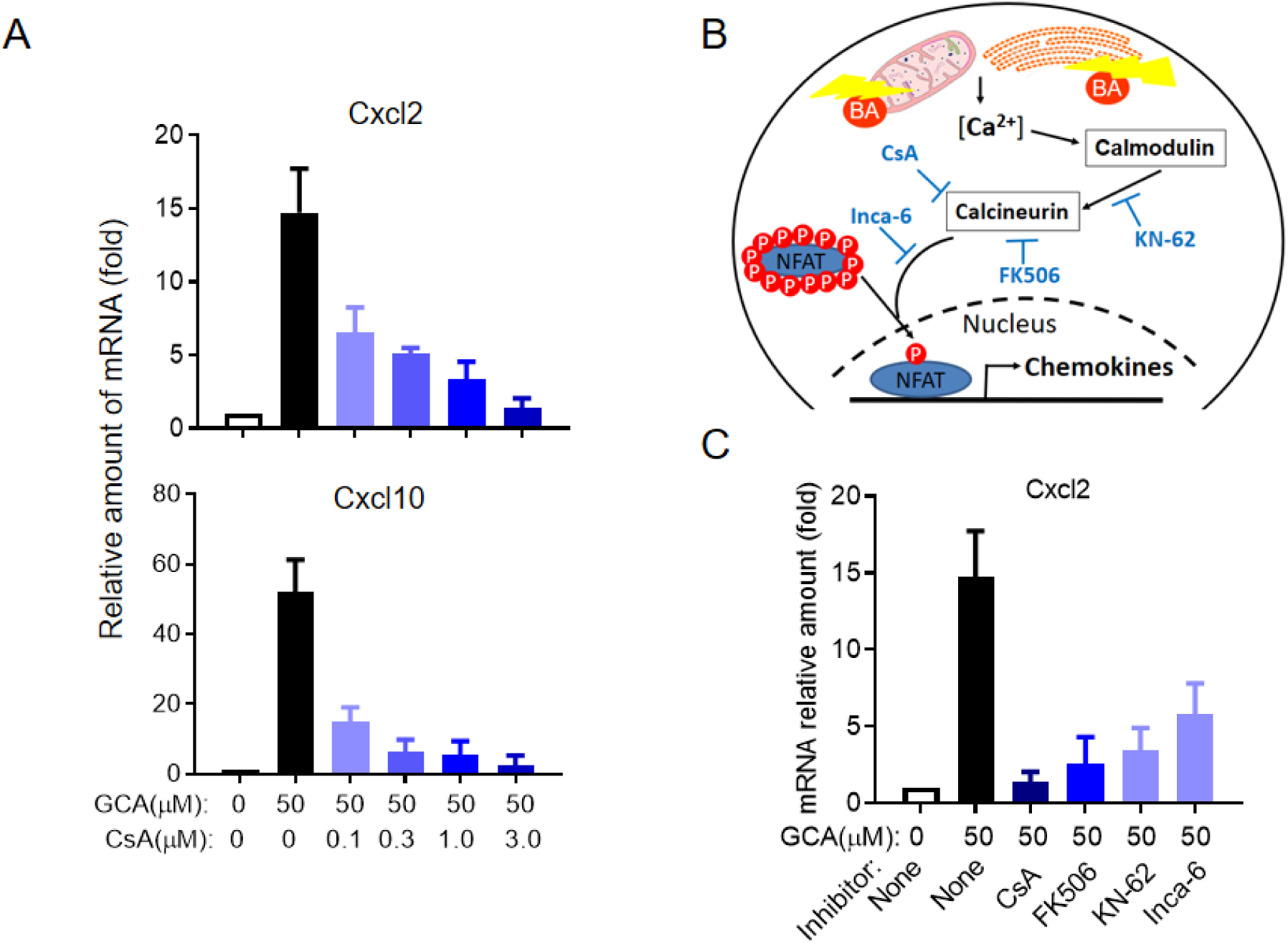
Inhibitors involved in Ca2+/NFAT signaling pathway repressed bile acid (BA) induction of inflammatory chemokines in mouse hepatocyte cultures. **A**, CsA repressed GCA induction of Cxcl2 and Cxcl10 mRNA expression in mouse hepatocytes (24 h treatment). **B**, A diagram of Ca^2+^/NFAT signaling pathway and inhibitors involved in NFAT activation. **C**, GCA induction of Cxcl2 expression were repressed by inhibitors involved in Ca^2+^/NFAT signaling in mouse hepatocytes. 24 h treatment, CsA (3 µM), FK506 (5 µM), KN-62 (3 µM), Inca-6 (40 µM). *p<0.05, n>=4). Data normalized to Gapdh, mean±SD, \**p*<0.05 n≥4.

### BA increased NFATc3 nuclear expression in mouse and human hepatocytes but not cholangiocytes

Nfatc1 and c3 are expressed in mouse cholangiocytes(20), but it is not known which isoforms are expressed in hepatocytes, nor whether BA alter their expression. To address these questions, we first analyzed the expression levels (mRNA) of the four different isoforms of NFATc/Nfatc in cultured primary hepatocytes from humans and mice. As shown in Figure 2A, NFATc1 and c3 are expressed in both human and mouse hepatocytes, whereas NFATc2 was also detected in human hepatocytes. Neither cells expressed significant amount of NFATc4. We also confirmed that Nfatc1 and c3 are expressed in mouse cholangiocytes as previously described (20) (Data not shown). We next determined whether BA altered the mRNA expression of Nfatc isoforms in mouse hepatocytes. 100 µM TCA treatment slightly but significantly increased the mRNA levels of Nfat isoform c1, c2 and c3 in these cells, where isoform c4 remained essentially undetectable (Fig 2B). Because NFATc3 is the most abundantly expressed isoform in both human and mouse hepatocytes, we next examined whether BA treatment led to Nfatc3 nuclear translocation in mouse hepatocytes. As shown in Figure 3A, at pathophysiological concentration (100 µM), TCA but not taurodeoxyursocholic acid (TUDCA) increased nuclear expression of Nfatc3 protein in these cells, where cytoplasmic expression of Nfatc3 protein was slightly decreased. Increased nuclear accumulation of NFATc3 protein was also found in primary human hepatocyte cultures when treated with GCDCA and TDCA, but not TUDCA (Fig.3B), indicating human and mouse share a common mechanism in hepatic NFAT activation. To examine whether other BA could cause NFAT nuclear translocation and also to test whether inhibitors of the Ca^2+^-calmodulin-calcineurin-NFAT signaling cascade could reduce BA-induced nuclear accumulation of NFAT, we treated mouse hepatocytes with CDCA, GCA and TDCA. As demonstrated in Figure 3C, GCA and TDCA treatment increased Nfatc3 nuclear expression, where CsA, KN-62 and FK506 each reduced GCA-induced Nfatc3 nuclear expression. Because Nfatc3 is also abundantly expressed in mouse cholangiocytes, we wondered whether BA could also induce its nuclear translocation in these cells. Western blot analysis did not show the same effects as seen in the hepatocyte cultures when even higher concentrations of TCA (400 µM) were applied (supplementary Figure S1), where no substantial induction of chemokine Ccl2, Cxcl2 and Cxcl10 were detected in these cells even when they were exposed to 1 mM TCA.Together, these findings indicate that the role of NFAT in BA induction of chemokines is an hepatocyte specific event in cholestatic livers.

**Figure 2.**
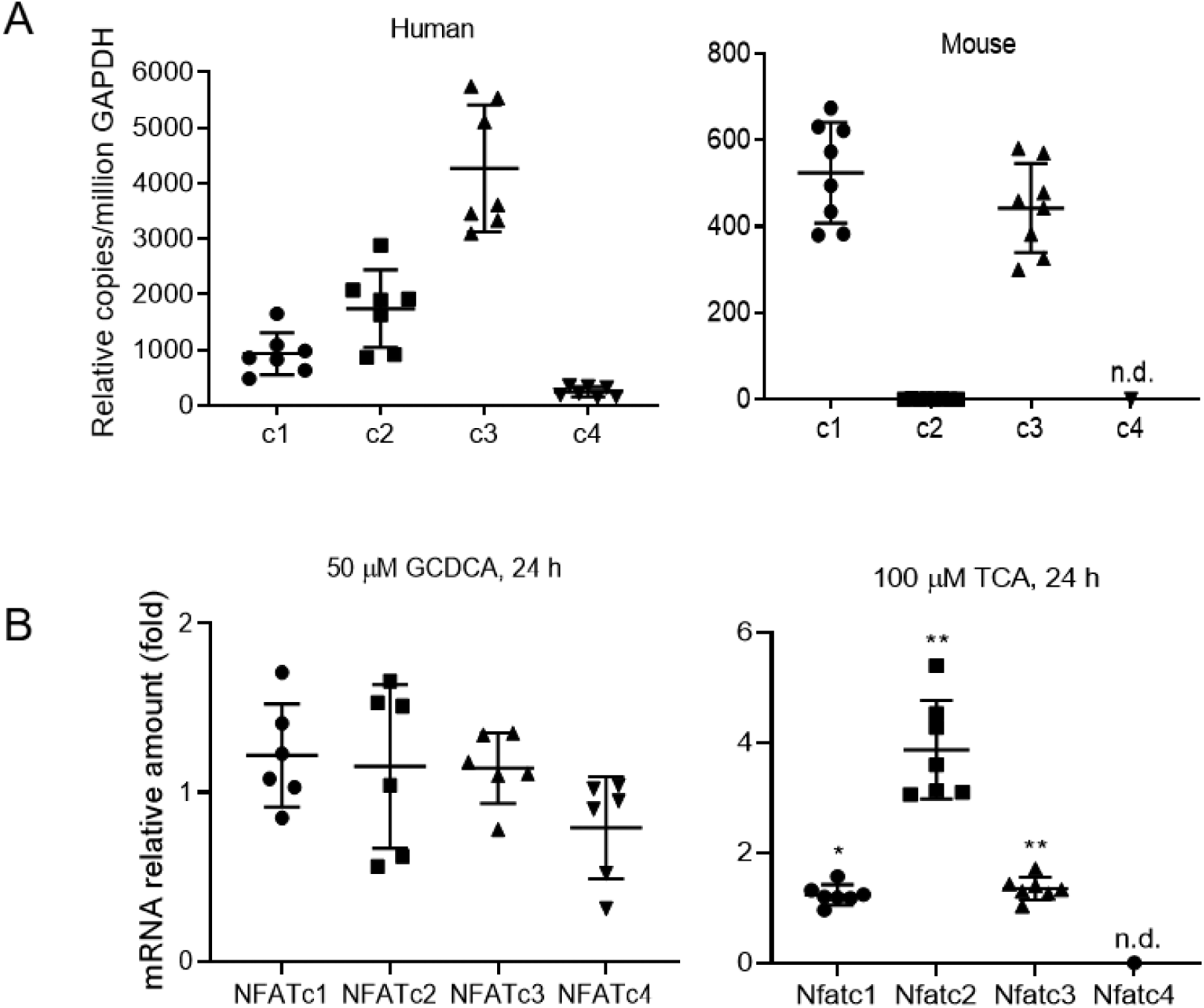
**A**, Message RNA expression of NFAT isoforms in human (left) and mouse (right) hepatocytes. Each dot represents an individual. **B**, Major endogenous bile acid significantly induced the mRNA expression of mouse but not human NFAT/Nfat isoforms c1, c2 and c3. The expression of each isoforms in control cells was set as 1. mean±SD, n=6-8, \**P*<0.05, \*\**P*<0.01, n.d., non-detectable.

**Figure 3.**
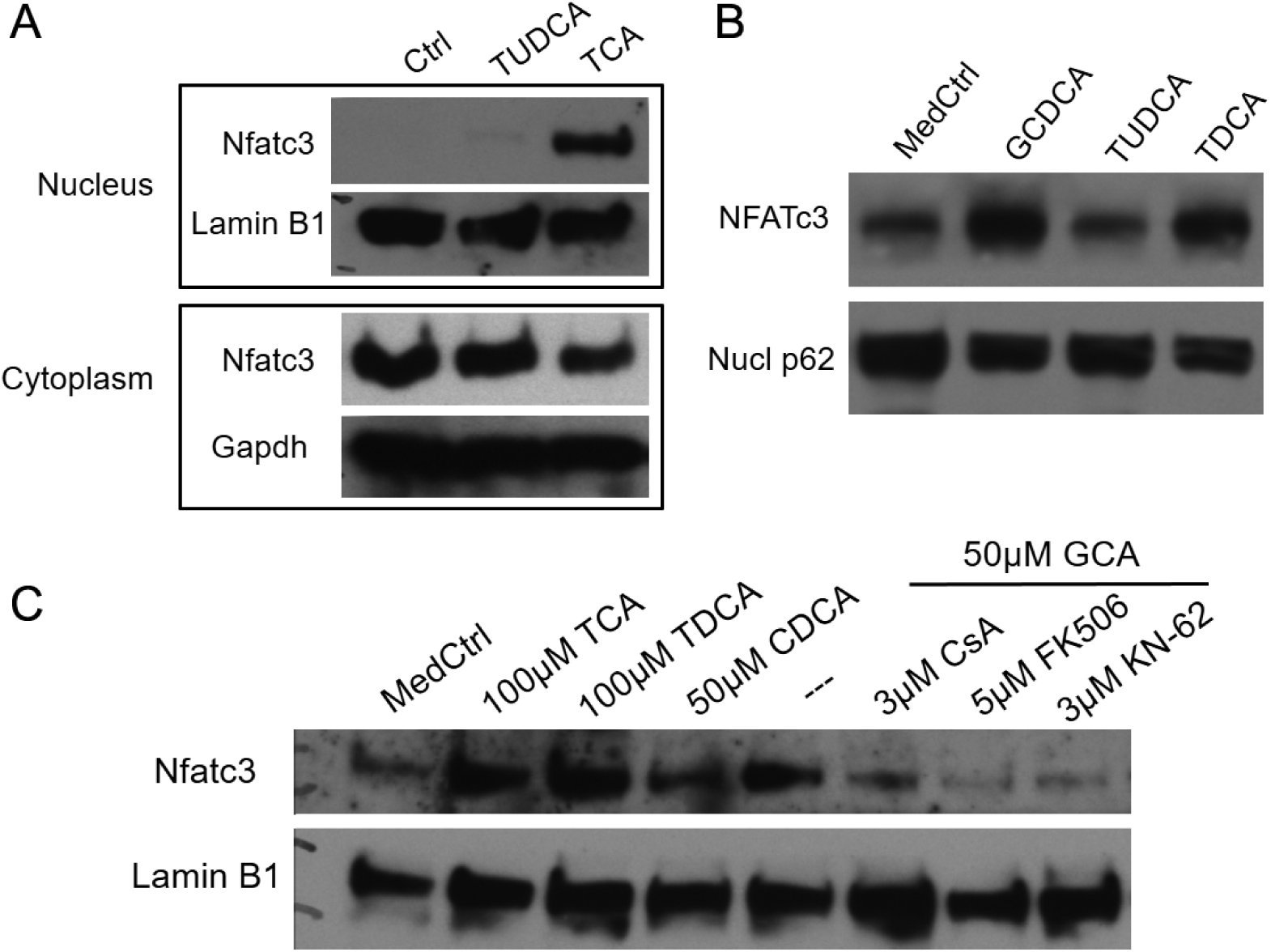
Western blots demonstrated that bile acids induced NFATc3/Nfatc3 nuclear translocation in both mouse and human hepatocytes. **A**. mouse hepatocytes were treated with TCA (100µM) or TUDCA (100µM) for 24 h. Lamin B1 as a nuclear protein marker, Gapdh as a cytoplasmic protein marker. **B**, nuclear protein from human hepatocytes exposed to GCDCA (100µM), TDCA (100µM) or TUDCA (100µM) for 24 h. Nucleoporin p62 (Nucl p62) as loading control. **C**, nuclear protein from mouse hepatocytes exposed to bile acids and also treated with inhibitors of Ca^2+^/NFAT signaling.

### Increased NFAT nuclear expression in cholestatic mouse livers was associated with elevated hepatic levels of inflammatory cytokines

To examine whether NFAT expression is altered in cholestatic livers, we first analyzed the expression (mRNA) of Nfatc isoforms in the livers of three cholestatic mouse models and compared with their corresponding experimental controls. These models and the degree of liver injury have been characterized in our previous reports (4, 18) and include the Abcb4^-/-^ mice (a model of sclerosing cholangitis), BDL mice (an obstructive cholestasis) and 1% cholic acid fed mice (mild cholestasis due to BA overload). Q-PCR analysis revealed that Nfatc4 was undetectable in the livers of all these mice. Both Nfatc1 and c3 were moderately expressed in the livers of these cholestatic animals at essentially the same level but neither were substantially altered. In contrast, the expression of Nfatc2 was significantly increased in the liver of all three cholestatic models (up to 6-fold in *Abcb4*^-/-^ mice) (Figure 4), although its basal level of expression was substantially lower (∼0.4%) than Nfatc1 and c3. To check whether Nfat protein expression had been altered in these cholestatic livers, we isolated cytoplasmic and nuclear fractions from liver tissue. Western blot detected significantly higher nuclear expression of Nfatc3 in both *Abcb4*^-/-^ livers and BDL livers than in their corresponding controls (Figure 4B). This increased nuclear expression of Nfatc3 was further supported by immunohistochemical staining of liver sections from *Abcb4*^-/-^ mice which showed increased staining throughout the liver lobules (Figure 4C). Furthermore, gene expression analysis revealed that expression (mRNA) of Cxcl2 was also significantly increased in the livers of all these cholestatic mice (Figure 4D).

**Figure 4.**
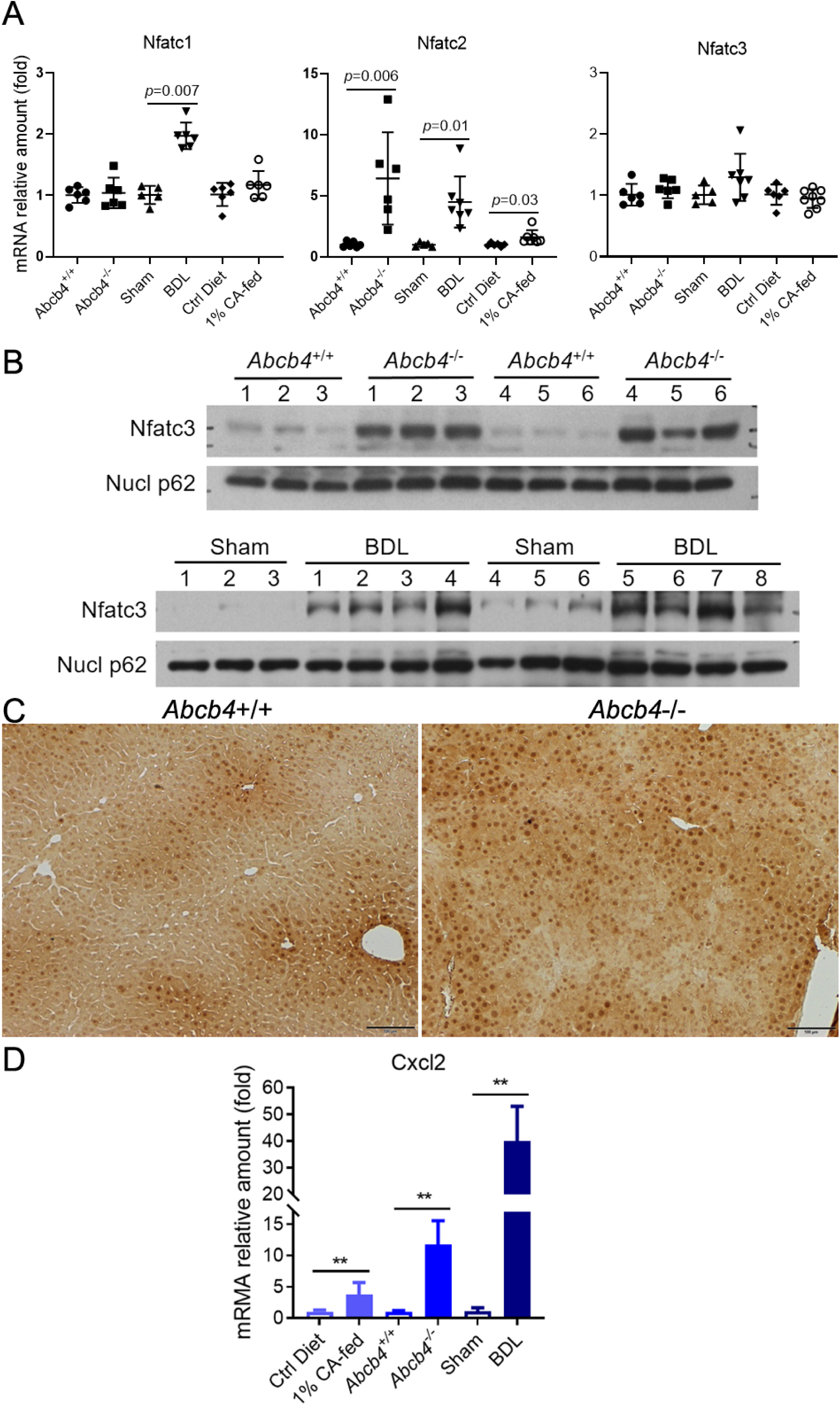
Elevated hepatic Cxcl2 expression was associated with increased nuclear protein expression of Nfatc3 in the livers of cholestatic mice. **A**, Nfatc2 mRNA expression was increased in cholestatic mouse livers. **B**, increased nuclear expression of Nfatc3 in *Abcb4*^-/-^ (top) and bile duct ligated (BDL) (bottom) mouse livers. **C**, immunohistochemical staining of Nfatc3 expression in the liver of wild-type and cholestatic *Abcb4*^-/-^ mice. **D**, hepatic Cxcl2 mRNA expression in *Abcb4*^-/-^, BDL, and CA-fed mice. 1% cholic acid (CA) fed for 7 days, 6-week old *Abcb4*^-/-^, and 7-day BDL. Nucl p62, nuclear protein p62 as nuclear protein loading marker. mean±SD, n=6-8, \*\**P*<0.01.

### Knockdown of Nfatc3 reduced mRNA expression of inflammatory genes in mouse hepatocytes

To determine whether Nfatc3 plays a direct role in BA induction of inflammatory genes in the liver, we knocked down Nfatc3 expression using siRNA transfection in mouse hepatocytes. As demonstrated in Figure 5, diminished Nfatc3 expression resulted in significant reduction of inflammatory genes at both basal level and after TCA stimulation, including Ccl2, Cxcl2, and Cxcl10, whereas the bile acid receptor Fxr mRNA was increased in these cells. Because a previous study (14) revealed a series of Nfatc2 targets in mouse T-cells using ChIP-Seq, including Ccl3, Cxcr4, Egr1, Egr3, Icam1, Vcam1, we also assayed the expression of these genes in Nfatc3 knockdown mouse hepatocytes. We found reduced expression of Egr1 and Icam1 (Figure 5). However, Vcam1 expression was not significantly changed, while Ccl3 and Egr3 were essentially undetectable (data not shown). Together, these results indicate that there are significant differences in NFAT targets between T-cells and hepatocytes and that NFAT plays an important role in BA induction of inflammatory genes in mouse hepatocytes.

**Figure 5.**
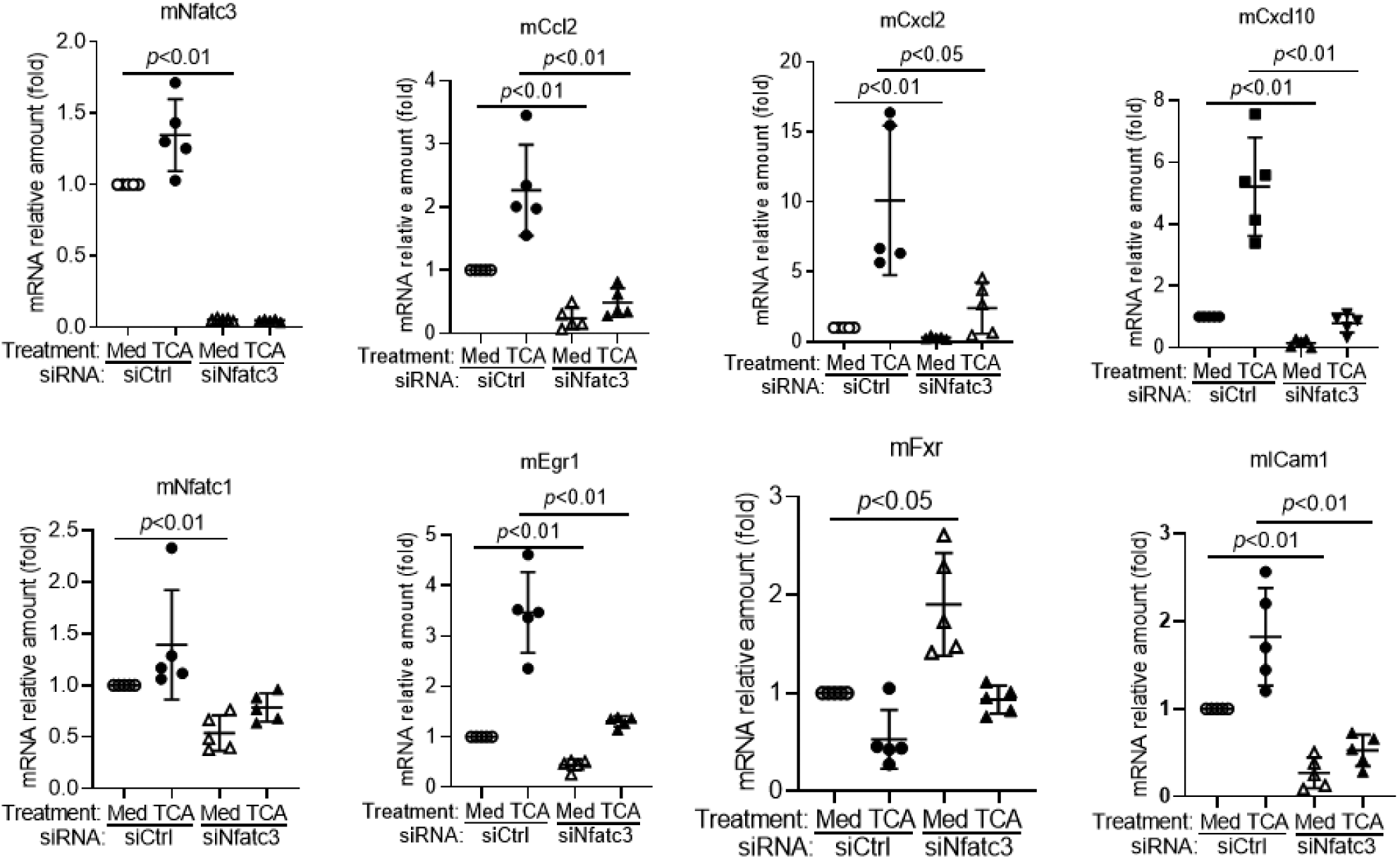
Knockdown of Nfatc3 reduced the expression of inflammatory genes in mouse hepatocyte cultures.

### Increased NFAT nuclear expression in the livers of patients with PBC and PSC is associated with elevated levels of IL-8

To verify whether altered hepatic expression of NFATc isoforms also occurs in human cholestasis, we first analyzed a few liver specimens from patients collected at Yale Liver Center. This includes 4 normal controls, 2 steatosis, 2 biliary atresia, 1 PBC and 1 PSC. Western blots revealed more NFATc3 protein in the nuclear fraction of livers from cholestatic patients but not from steatotic donors, whereas there were no substantial differences in the cytoplasmic fraction among these donors (Figure S2). To validate these preliminary observations we analyzed an additional group of liver specimens from cholestatic patients with PBC and PSC collected by the Liver Tissue Cell Distribution System in University of Minnesota. As demonstrated in Figure 6, the nuclear expression of NFATc3 protein was significantly increased in the livers of both PBC (3.8-fold) and PSC (3.4-fold) patients when compared to the non-diseased control livers while their cytosolic protein levels did not differ significantly among these three groups. We also detected significant increases in mRNA of IL-8 in PBC and PSC livers (29 and 43-fold, respectively) when compared to normal liver tissue controls (Figure 6C). Further data analysis of IL-8 expression and the amount of NFATc3 nuclear protein in PBC livers indicated that the nuclear but not cytosolic expression of NFATc3 protein correlated directly with hepatic IL-8 mRNA levels (Fig 6D. r=0.663, *p*=0.005, n=16), suggesting that NFATc3 up-regulates IL-8 expression in these cholestatic livers. To examine whether there are any differences at mRNA level in NFATc isoforms between healthy livers and cholestatic livers, we analyzed their expression by Q-PCR. In contrast to the cholestatic mice, mRNA levels for the four isoforms did not differ significantly between disease groups and the healthy controls (supplementary Fig.S3). However, at the basal level, the relative mRNA abundance of these four isoforms in the healthy control livers was NFATc1 = NFATc3 > NFATc2 >> NFATc4 (data not shown), similar to what was seen in mouse livers.

**Figure 6.**
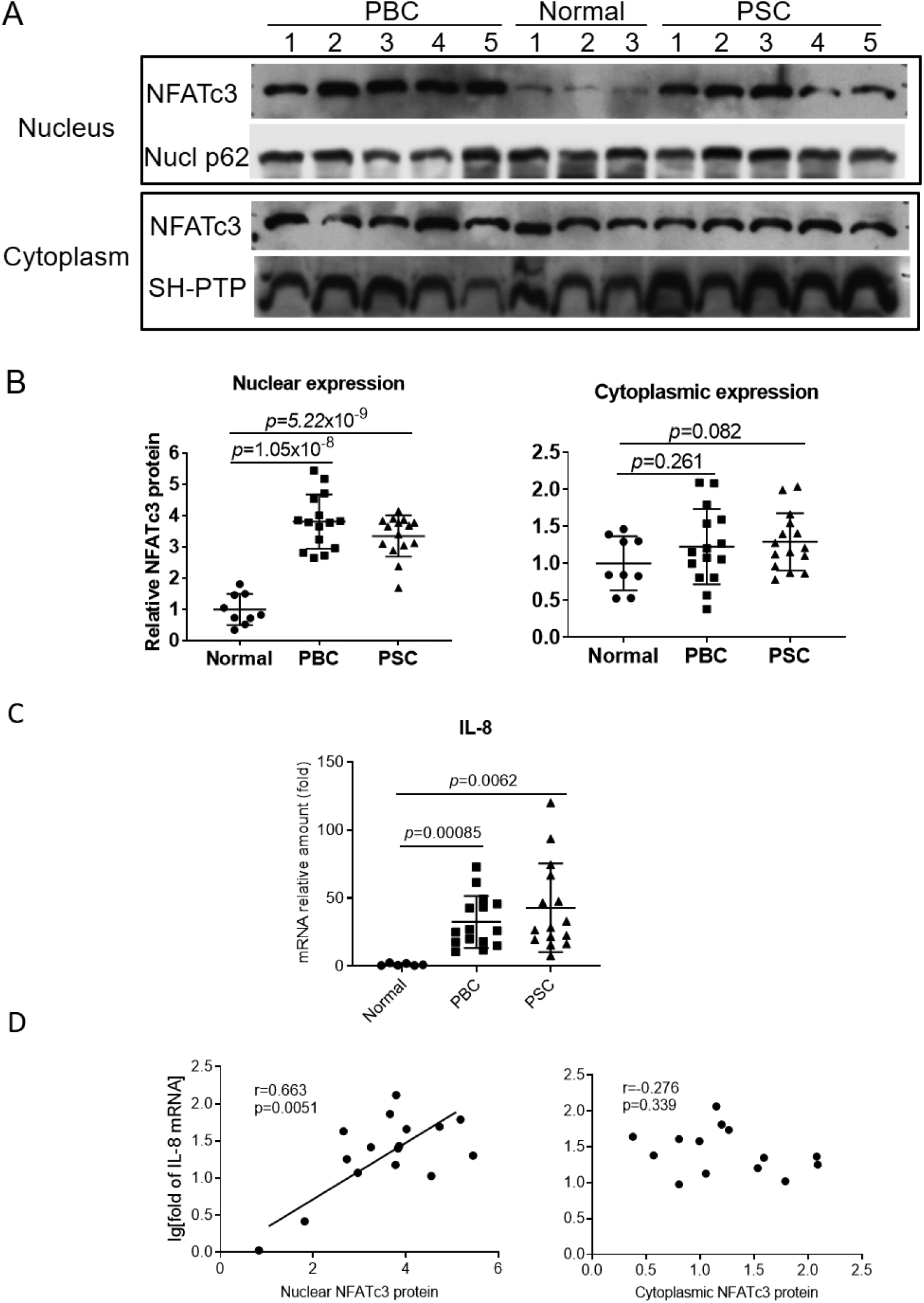
**A**, Western blot of nuclear and cytoplasmic expression of NFATc3 protein in patients with PBC, PSC and normal livers; **B**, Relative amount of NFATc3 protein by densitometry analysis. Nucl p62 and SH-PTP are loading controls for nuclear and cytoplasmic protein, respectively). **C**, mRNA expression of IL-8 in the liver of patients with PBC and PSC. **D**, Correlation analysis of the hepatic levels of nuclear NFATc3 protein and IL-8 mRNA in PBC patients.

### BA stimulated IL-8 expression through its NFAT response element

Our previous studies indicated that cholestatic levels of BA-stimulated IL-8 mRNA expression in primary human hepatocyte cultures(4). To test whether NFAT mediates this stimulation, we generated a human IL-8 promoter reporter construct that contains an NFAT response element ((21) and Figure 7A). When this reporter construct was transfected into Huh7-BAT cells (a cell line that is stably transfected and expresses the bile acid uptake transporter NTCP/SLC10A1 (19)), it demonstrated strong basal activity, ∼22-fold compared to the control vector pGL3-basic (Figure 7B). When the transfected cells were treated with BA (50 µM), GCDCA, GCA and TCA all significant increased IL-8 promoter activity, whereas unconjugated cholic acid did not demonstrated the same effect (Fig.7B). To examine whether the conjugated BA induction of the IL-8 promoter is NFAT dependent, we mutated its NFAT response element. As shown in Figure 7B, mutation of this NFAT response element greatly reduced both the basal activity and the BA inducibility of the IL-8 promoter. To further confirm NFAT’s role in BA-stimulated IL-8 expression, we performed ChIP-PCR in Huh7-BAT cells. Because GCA and TCA greatly stimulated IL-8 promoter activity in reporter assays (Fig.7B), we treated these cells with 50 µM GCA or TCA for 4 hr. Figure 7C demonstrated increased amounts of IL-8 promoter DNA in GCA and TCA treated cells than in the control cells. Together, these findings indicate that endogenous conjugated BA stimulate human IL-8 hepatic expression by activating NFAT.

**Figure 7.**
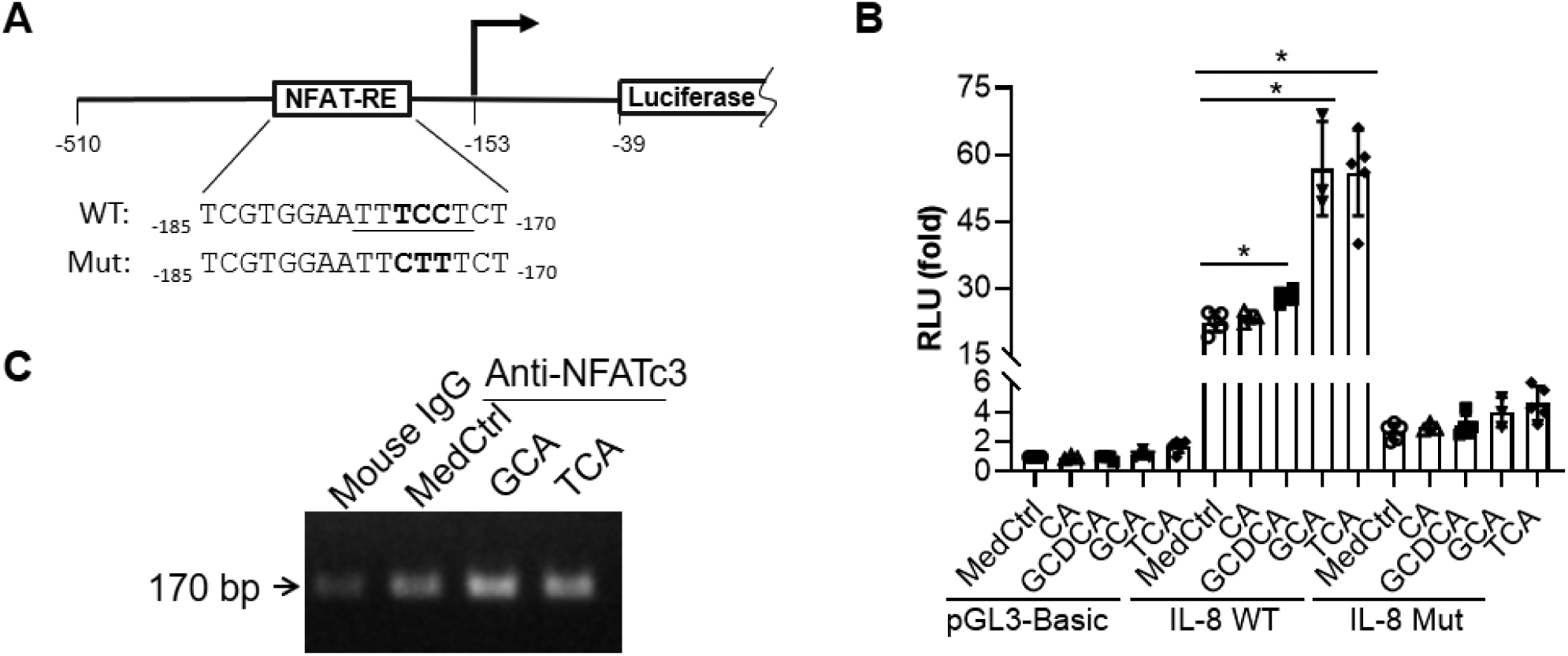
Bile acids stimulate human IL-8 promoter activity through a NFAT response element (NFAT-RE) in gene reporter assay and ChIP-PCR. **A**, a diagram of human IL-8 promoter reporter constructs in pGL3-basic vector. **B**, Relative promoter activities of human IL-8 in transfected Huh7-BAT cells (a cell line that stably expresses NTCP) from dual-luciferase reporter assay. **C**, Agarose gel electrophoresis showed that bile acids treatment increased NFATc3 binding to IL-8 promoter in Huh7-BAT cells using ChIP-PCR assay. Of note, for the reporter assay, 24 hr after transfection, cells were treated with 50 μM of the indicated bile acids for 16 h. Data was normalized to Renilla luciferase activity, and pGL3-basic activity in control medium (MedCtrl) was set as 1. For ChIP-PCR, cells were treated with 50 μM GCA or TCA for 4 hr. CA, cholic acid; GCDCA, glycochenodeoxycholic acid; GCA, glycocholic acid; TCA, taurocholic acid. mean±SD, n≥3, \**p*<0.05.

## Discussion

In this report, we examined the functional role of NFAT in cholestatic liver injury. We found that: 1) NFAT isoforms are expressed in hepatocytes from humans and mice (Fig. 2); 2) Under cholestatic conditions, BA-stimulated expression of inflammatory cytokines in mouse and human hepatocytes was associated with the activation of transcription factor NFAT (Figs. 1 & 3). Blocking this activation by using pathway specific inhibitors or knocking down Nfatc3 expression reduced BA induction of these proinflammatory genes in mouse hepatocytes (Figs. 1 & 5); 3) Increased expression of NFATc3/Nfatc3 nuclear protein was found in cholestatic livers from both patients with PBC and PSC and in mice with *Abcb4*-deficiency or after BDL where elevated levels of inflammatory cytokines were also found (Figs. 4 & 6); Finally, 4) both gene reporter assays and ChIP-PCR demonstrated that BA-induced human IL-8 expression is mediated through NFAT response elements in its proximal promoter (Fig. 7). Based on these observations, we conclude that NFAT plays an important role in the pathogenesis of cholestatic liver injury, a novel mechanism of BA initiated hepatic inflammatory response.

Studies in T-cells and other immune cells have established that NFATc isoforms are activated through the Ca^2+^/calmodulin/Calcineurin signaling pathway. When there is a sustained elevation of intracellular Ca^2+^, the NFATc isoform translocates to the nucleus and acts as a transcription factors to modulate gene expression (7, 22, 23). There is a substantial literature indicating that BA increase cytosolic Ca^2+^ levels in hepatocytes (24-26). While those studies mainly demonstrate transient changes in cytosolic calcium, it seems likely that more chronic elevations could occur. Both ER and mitochondria play a pivotal role in regulating Ca^2+^ signaling, and BA also cause ER-stress and mitochondrial damage when they reach cholestatic levels (27-29). Thus, it is likely that NFATc is activated in cholestatic hepatocytes since NFATc isoforms are expressed in these cells (Fig. 2). In addition, Nfatc3 protein accumulates in the nucleus of BA treated mouse and human hepatocytes (Fig. 3) but not in the livers of patients with other forms of liver injury, e.g. steatosis (Figure S2), further supporting the involvement of hepatic NFATc as a contributor to the cholestatic inflammatory response. Of note, NFATc3 is also present in the nuclei of hepatocytes under normal conditions as detected by both Western blot and IHC (Figs.3 and 4B), suggesting that hepatic NFATc3 may also play a role in maintaining normal liver physiology. Incomplete regeneration of the liver after partial hepatectomy seen in *Nfatc3*^-/-^ mice would support this view (30). However, during cholestasis, nuclear NFATc3 expression increases in response to BA stress, resulting in stimulating the expression of inflammatory genes. This also seems to be hepatocyte specific in the liver, because BA did not increase the expression of nuclear Nfatc3 or chemokines in cholangiocyte cultures (Supplementary Fig.S1 and (4)), nor did we observe increased Nfatc3 nuclear expression in the bile ducts of *Abcb4*^-/-^ livers (Fig.4B). Since there is little if any expression of NFATc4 in the liver (Fig.2), we conclude that this isoform does not play a role in cholestatic liver injury. However, although the level of NFATc2 was substantially lower than NFATc3 and NFATc1, both NFATc1 and NFATc2 are detected in the liver (Figs. 2, and S2 and (20)), so it remains to be determined whether they play any role in BA-stressed cholestatic hepatocytes, given that NFATc isoforms modulate gene expression in both a stimulus- and a cell type–specific manner. Future studies may address these questions.

Activation of hepatic NFATc3 is at least partially responsible for BA induction of inflammatory genes because inhibitors blocking Nfatc3 nuclear translocation or knockdown of Nfatc3 significantly reduced BA stimulation of inflammatory gene expression in mouse hepatocytes(Figs. 1, 3 & 5). The response elements for NFAT in many inflammatory genes have been identified by ChIP-Seq in mouse T-cells, including Icam1 and Egr1, supporting NFAT’s role in regulating the immune response (14). Most importantly, we demonstrated that BA-stimulated IL-8 expression was mediated through NFAT response element in the human IL-8 promoter, as demonstrated by reporter assay and ChIP-PCR assay in Huh7-BAT cells (Fig.7), and consistent with elevated levels of IL-8 in the livers of patients with PBC and PSC (Fig.6). Because our findings indicate that NFAT plays an important role in BA-stimulation of inflammatory cytokine expression, and because the inflammatory response plays a critical role in the pathogenesis of cholestatic liver injury(31, 32), one may speculate that blocking hepatic NFAT activation would reduce liver injury in cholestasis. Future studies should test this hypothesis. Interestingly, a CsA analog, NIM811, has previously been shown to reduce liver injury in the BDL cholestatic mouse (33) and we found that NIM811 had similar effects as CsA in suppressing BA-induction of cytokines in mouse hepatocytes (Data not shown).

Previous reports indicate that NFAT signaling cross talks with the innate immune response via TLR signaling(34, 35). Our prior study revealed that Tlr9 was involved in the BA-induced hepatic inflammatory response (4), while a recent report indicates that Ca^2+^/NFAT signaling can be down-stream of Tlr9 activation (36). In that report, activated Tlr9 stimulates Burton’s tyrosine kinase (BTK) activity, resulting in phosphorylation and activation of phospholipase Cγ. The activated phospholipase Cγ then initiates Ca^2+^/NFAT signaling. The involvement of Ca^2+^/NFAT signaling was also seen in TLR9-induced IL-10 secretion in human B-cells where Burton’s tyrosine kinase also played a role (37). Thus, it remains to be determined whether BA-activated Tlr9 and Ca^2+^/NFAT signaling are two independent signaling pathways or are part of the same cascade of events in cholestatic hepatocytes. If the latter, Ca^2+^/NFAT signaling would be down-stream of TLR9. Blocking NFAT signaling would also stop TLR9 activation, suggesting that NFAT is the master controller in BA induction of proinflammatory genes in hepatocytes. Future studies may address this question.

In summary, our findings indicate that the transcription factor NFAT plays an important role in regulating the expression of inflammatory genes in cholestatic hepatocytes, a novel mechanism in the inflammatory response induced by BA. These findings suggest that the NFAT signaling pathway should provide new targets for treating cholestatic disorders.

## Acknowledgements

We are grateful to Kathy Harry and Maoxu Ge for their excellent technical support. We also thank Drs. Mario Strazzabosco and Romina Fiorotto here in our section for providing primary mouse cholangiocytes, Dr. Gregory Gores, Mayo Clinic, Rochester, Minnesota for sharing Huh7-BAT cells.

## Notes

Grant Support: This study was supported by National Institutes of Health Grants DK34989 (Yale Liver Center), DK25636 (to J.L.B.).

**Disclosures:** The authors have declared that no conflict of interest exists.

