## Supplementary data for "Hepatic NFAT signaling regulates the expression of inflammatory cytokines in cholestasis"

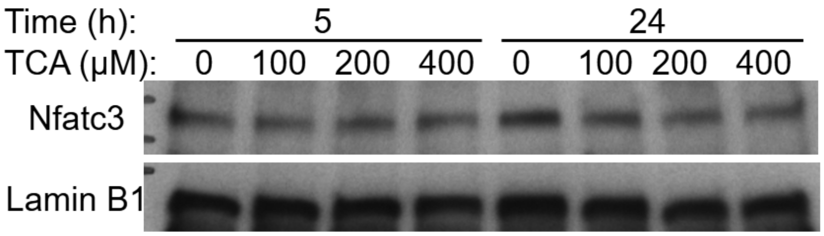


**Figure S1** Western blot analysis showed that taurocholic acid (TCA) did not cause Nfatc3 nuclear translocation in primary mouse cholangiocyte cultures. Cells were treated with indicated concentration of TCA and time.

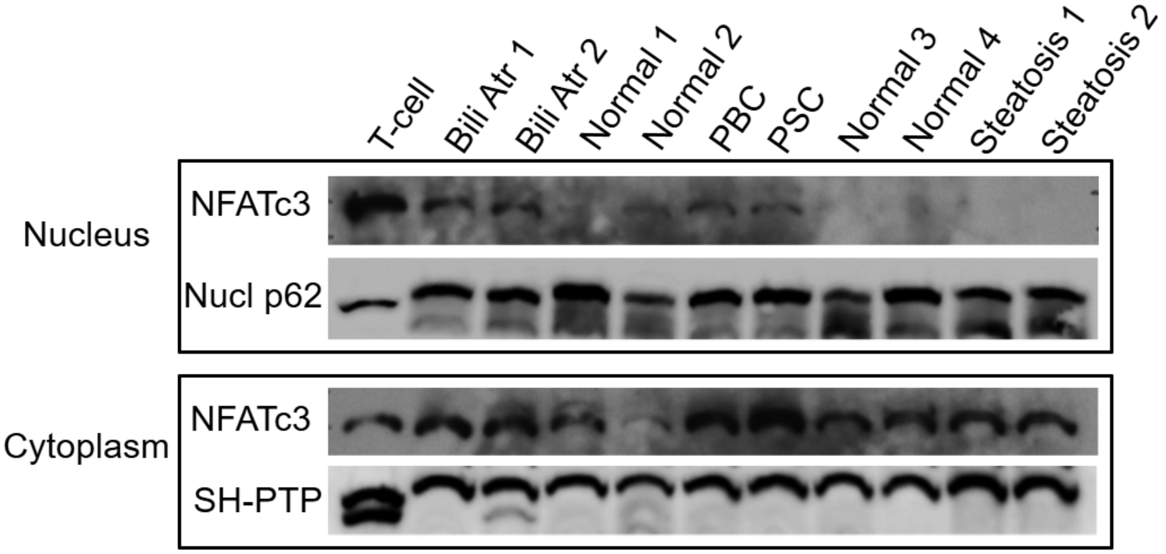


**Figure S2** Western blot demonstrates that increased nuclear expression of NFATc3 in the liver of cholestatic patients. Total T-cell lysate as positive control. Nucleoporin p62 (Nucl p62) as a nuclear marker, SH-PTP as a cytoplasmic loading control.Bili Atr, biliary atresia; PBC, primary biliary cholangitis; PSC, primary sclerosing cholangitis.

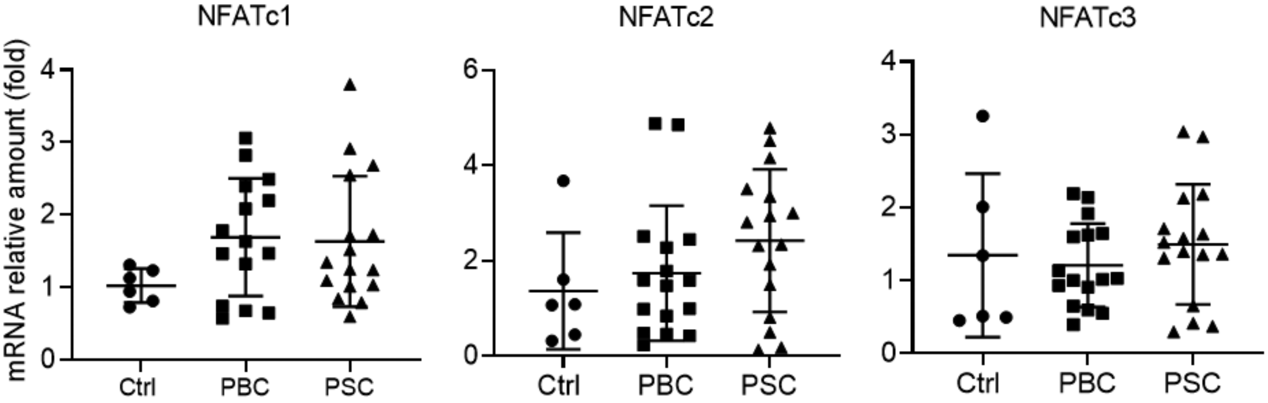


**Figure S3** Relative amount of NFAT isoform mRNA in the liver of healthy controls and cholestatic patients. Ctrl, normal healthy donor (n=6); PBC, primary biliary cholangitis (n=15); PSC, primary sclerosing cholangitis (n=15). Data normalized to GAPDH, mena±SD.
